# Programmed Switch in The Mitochondrial Degradation Pathways During Human Retinal Ganglion Cell Differentiation from Stem Cells is Critical for RGC Survival

**DOI:** 10.1101/638585

**Authors:** Arupratan Das, Claire M. Bell, Cynthia A. Berlinicke, Nicholas Marsh-Armstrong, Donald J. Zack

**Affiliations:** Department of Ophthalmology, Wilmer Eye Institute, Johns Hopkins University School of Medicine, Baltimore, Maryland, USA; McKusick-Nathans Institute of Genetic Medicine, Johns Hopkins University School of Medicine, Baltimore, Maryland, USA; Department of Ophthalmology and Vision Science, University of California, Davis, CA, USA; Department of Molecular Biology and Genetics, Johns Hopkins University School of Medicine, Baltimore, Maryland, USA; The Solomon H. Snyder Department of Neuroscience, Johns Hopkins University School of Medicine, Baltimore, Maryland, USA; Institute of Genetic Medicine, Johns Hopkins University School of Medicine, Baltimore, Maryland, USA

**Keywords:** Stem cells, Human retinal ganglion cells (hRGCs), Glaucoma, Neurodegeneration, Autophagy-lysosome, UPS

## Abstract

Retinal ganglion cell (RGC) degeneration is the root cause for vision loss in glaucoma as well as in other forms of optic neuropathies. Genetic analysis indicated abnormal mitochondrial quality control (MQC) as a major risk factor for optic neuropathies. However, nothing is known on how MQC regulates human retinal ganglion cell (hRGC) health and survival. Human pluripotent stem cells (hPSCs) provide opportunity to differentiate hRGCs and understand the abnormal MQC associated hRGC degeneration in great detail. Degradation of damaged mitochondria is a very critical step of MQC, here we have used stem cell derived hRGCs to understand the damaged mitochondrial degradation pathways for hRGC survival. Using pharmacological methods, we have investigated the role of the proteasomal and endo-lysosomal pathways in degrading damaged mitochondria in hRGCs and their precursor stem cells. We find that upon mitochondrial damage with the proton uncoupler carbonyl cyanide m-chlorophenyl hydrazone (CCCP), hRGCs more efficiently degraded mitochondria than their precursor stem cells. We further identified that for degrading damaged mitochondria, stem cells predominantly use the ubiquitine-proteasome system (UPS) while hRGCs use the endo-lysosomal pathway. UPS inhibition causes apoptosis in stem cells, while hRGC viability is dependent on the endo-lysosomal pathway but not on the UPS pathway. This suggests manipulation of the endo-lysosomal pathway could be therapeutically relevant for RGC protection in treating glaucoma. Endo-lysosome dependent cell survival is also conserved for other human neurons as differentiated human cerebral cortical neurons also degenerated upon endo-lysosomal inhibition but not for the proteasome inhibition.

**SIGNIFICANCE STATEMENT:** Using human stem cells we have shown a switch in the mitochondrial degradation pathway during hRGC differentiation where endo-lysosomal pathway becomes the predominant pathway for cellular homeostasis and hRGC survival which is also true for human cortical neurons. These findings suggest manipulation of the endo-lysosomal pathway could be therapeutically relevant for RGC protection in treating glaucoma as well as for other neurodegenerative diseases.

## INTRODUCTION

Optic neuropathies such as glaucoma, Leber’s hereditary optic neuropathy (LHON), dominant optic atrophy (DOA) [1] and several other neurodegenerative diseases are associated with abnormal mitochondrial quality control (MQC) [2,3]. In almost all of these optic neuropathies, irreversible damage of retinal ganglion cells (RGCs) leads to complete blindness [1]. MQC involves mitochondrial dynamics, biogenesis and degradation. While each step of MQC is important for mitochondrial homeostasis, defects in mitochondrial degradation are particularly severe, as they result in an accumulation of damaged mitochondria and ultimately lead to cell death through apoptosis [4–6]. Macroautophagy is a conserved catabolic process in which damaged proteins or organelles are degraded through forming a double membrane structure around them with complex protein interactions known as autophagosomes followed by fusion with the lysosomes where the damaged materials are degraded [7–9]. Selective degradation of damaged mitochondria through the lysosome-mediated autophagic pathway is called mitophagy [10,11]. Apart from cell autonomous mitophagy, recent report has also shown that RGCs shed mitochondria at the mice optic nerve head (ONH) by the adjacent astrocytes, a process referred to as transmitophagy [12].

Investigating human RGC-specific mitochondrial degradation pathways at the cellular level has been challenging due to the unavailability of hRGCs. Although studies in rodent models using both in-vivo and purified primary RGCs have given great insights into the molecular pathways involved in RGC survival [13–18], attempts to implement this knowledge in treating human optic neuropathies have been largely unsuccessful due to the inherent differences between rodent and human RGCs [19]. Therefore, in order to successfully move forward it is essential to have human stem cell-derived RGCs which will enable us to have a comprehensive understanding of MQC and its potential role in hRGC survival. It will further enable us to study the adaption of the MQC pathways during the course of RGC differentiation by comparing the process both in the stem cells and in differentiated RGCs.

Healthy mitochondrial homoeostasis in adult human stem cells is required to prevent stem cell aging and maintaining pluripotency [20]. The endo-lysosomal and proteasomal pathways are the two major cellular quality control pathways for clearing damaged organelles and proteins. However, it is unclear how hRGCs and their origin stem cells use either pathway for maintaining mitochondrial homeostasis. Studies in mice have shown mitophagy is required for the self-renewal [21,22] and differentiation [23] of hematopoietic stem cells (HSCs) as well as for cancer stem cell maintenance [24,25] in humans. The ubiquitin proteasome system (UPS) is highly active in hPSCs and upon cellular differentiation, the proteasome remains active but at a reduced level [26,27]. It is still unclear if hPSCs use the UPS system for degrading damaged mitochondria.

Several studies in mice have shown programmed mitophagy is required for RGC differentiation [28,29], and an *E50K* mutation in the autophagy adaptor protein optineurin (OPTN) has been shown to cause mitochondrial accumulation and RGC death [30]. Additionally, *OPTN^E50K^* mutation was also found in the severe form of normal-tension glaucoma (NTG) patients [31]. It is well accepted that the mitochondrial dynamics and quality control are central to mouse RGC viability [32]; however, the role of the lysosomal-autophagy and proteasomal pathways in degrading damaged mitochondria in hRGCs and its effect on hRGC survival are not yet understood.

In this study, we used small molecule-based hPSC differentiation and bead-based immunopurification to obtain highly pure, well-characterized hRGCs [33]. These hRGCs were used to investigate the role of the endo-lysosomal and the proteasomal pathways in clearing damaged mitochondria in comparison to their precursor stem cells. Our study shows hRGCs predominantly use the endo-lysosomal pathway for degrading damaged mitochondria to prevent apoptosis, whereas hPSCs primarily use the proteasomal pathway for mitochondrial clearance and cell survival.

## MATERIALS AND METHODS

### Reporter line Generation

H9 (WiCell, Madison, https://www.wicell.org) human embryonic stem cells with the BRN3B-P2A-tdTomato-P2A-THY1.2 reporter were developed in our lab [33] and used in this study as H9-ESCs. iPSCs (EP1) were developed in our lab [34] and used here as EP1-iPSCs. EP1 with the BRN3B-P2A-tdTomato-P2A-THY1.2 reporter was made by CRISPR/Cas9-based gene editing using a gRNA plasmid with Cas9 and puromycin selection, and the donor plasmid with the reporter genes as used before [33]. In Brief, EP1-iPSCs were transfected with the DNA-In stem reagent (MTI-GlobalStem/Thermo Fisher) following fresh media change after 24 hrs. After 40 hours of transfection, cells were selected against puromycin (0.9 μg/ml) for 24 hrs and recovered for 4-5 days with fresh media without puromycin. To isolate positive clones, cells were re-plated at clonal density, and single colonies were genotyped by PCR as described in the previous study [33].

### Human RGC Differentiation and Immunopurification

RGC reporter lines were plated on 1% (vol/vol) Matrigel-GFR (BD Biosciences) coated dishes and differentiated using small molecules as described in the previous study [33]. Successful RGC differentiation was monitored by tdTomato expression and purified during day 40-45 after dissociation with Accumax cell dissociation solution (Innovative Cell Technologies), as described in the previous study [33].

### hPSC and RGC Maintenance and Drug Treatments

hPSCs and RGCs were cultured and maintained on 1% (vol/vol) Matrigel-GFR (BD Biosciences) coated dishes in mTeSR and N2B27 media [33] respectively. Stem cells and RGCs were cultured in 37°C hypoxia (10% CO_2_, 5% O_2_) and normoxia (5% CO_2_) incubators, respectively. The following drugs were used in this study: CCCP (Sigma, # C2759), bafilomycin A1 (Sigma, # B1793), hydroxychloroquine (Fisher Scientific, # AC263010250), bortezomib (Selleckchem, # S1013), oligomycin (Millipore, # 495455), antimycin (Sigma, #A8674), oligomycin-antimycin (OA) drug combination used at 10μM and 4μM concentrations respectively, and MG132 (Millipore-Sigma, # M8699).

### Mitochondrial DNA Quantification by qPCR

After purification, RGCs were plated on Matrigel-coated tissue culture plates and grown for three days prior to the indicated drug treatments. Cells were dissociated using Accumax for 15min and quenched with the N2B27 media followed by centrifugation at 300Xg for 5min. DNA was isolated from the cell pellets using DNeasy Blood and Tissue kit (Qiagen) followed by simultaneous quantification of the mitochondrial and nuclear DNA content within the same sample using Taqman chemistry (Thermo Fisher) with StepOnePlus Real-Time PCR system (Applied Biosystems). Human mitochondrial DNA was detected via measurement of the very stable region on the mitochondrial ND1 gene [35] using following primers [36]; forward: 5’ CCTTCGCTGACGCCATAAA3’, reverse: 5’TGGTAGATGTGGCGGGTTTT3’, ND1-probe: 6FAM-5’TCTTCACCAAAGAGCC3’-MGBNFQ (6FAM and the MGBNFQ are the fluorescence reporter and quencher respectively). For an internal control, nuclear DNA content was measured using the human RNase P gene (TaqMan Copy Number Reference assay Catalog # 4403326).

For hPSCs (H9-ESCs/EP1-iPSCs), 15,000 cells were plated on each well of a Matrigel-coated 96-well dish. After 24 hrs of recovery, cells were treated with the indicated drugs and dissociated with Accutase (Millipore-Sigma) and quenched with mTeSR media (STEMCELL Technologies) containing blebbistatin (Sigma), followed by centrifugation at 300Xg for 5min to pellet the cells. Mitochondrial content for each sample was measured as explained above.

### Mitochondrial DNA Quantification by Flow cytometry

Flow cytometry-based measurements were done using mitochondria specific dye, mito tracker deep red (MTDR, Molecular probes) using cell sorter SH800 (Sony) on analyzer mode. 10,000-20,000 cells were analyzed at the FL-4 channel (far-red) to measure MTDR intensity. For 3 hr CCCP treatments (Fig. 1G, I), cells were labelled first while for 24 hrs treatments, cells were labelled after the drug treatments with media containing 10nM MTDR for 15min at 37°C. Cells were dissociated and centrifuged as explained in the qPCR method followed by suspension in media without MTDR for flow analysis.

**Figure 1.**
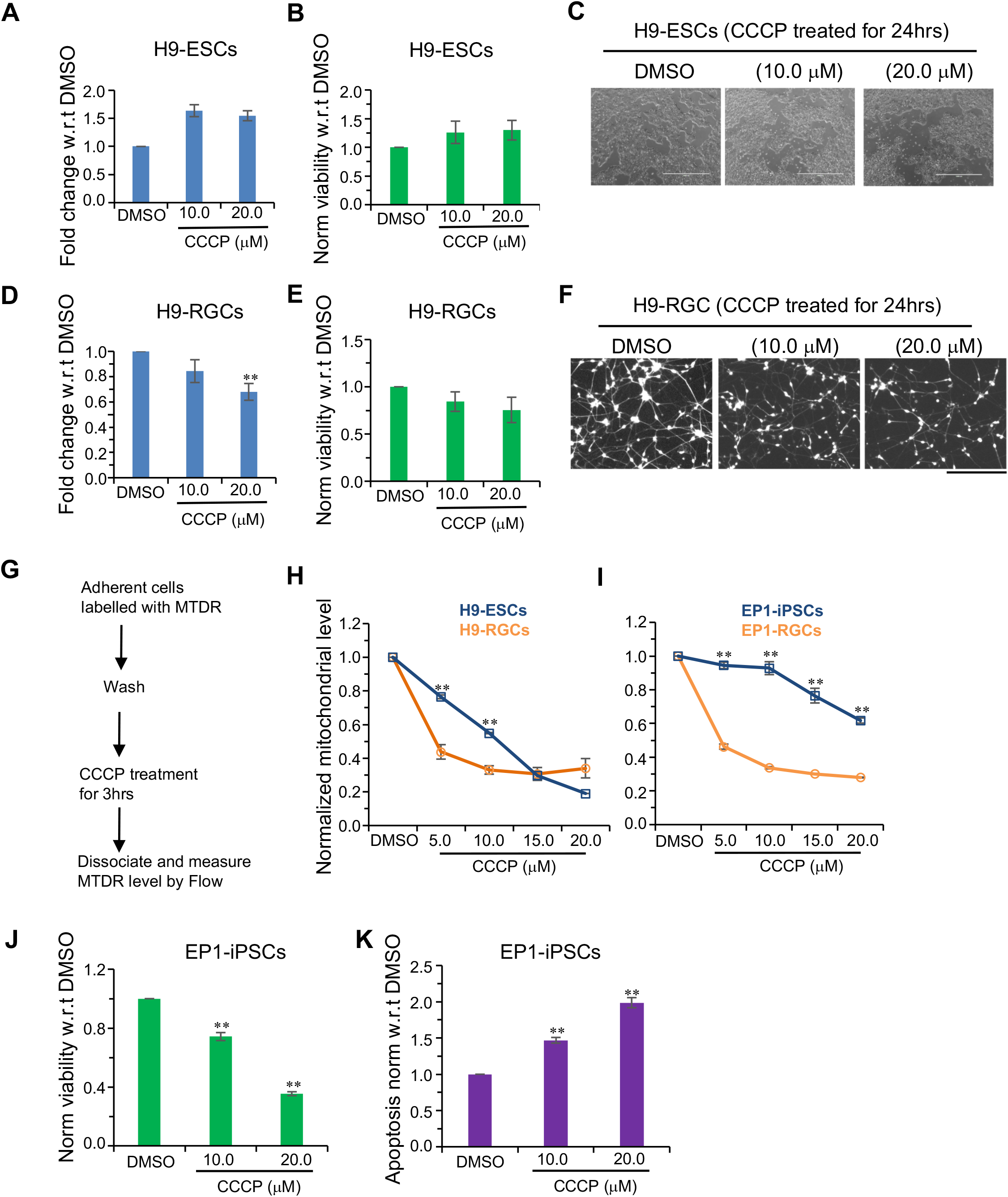
Mitochondrial degradation in stem cells and hRGCs upon CCCP treatment. (**A, D**) Mitochondrial content analyzed by qPCR for the mitochondrial gene ND1 and normalized with-respect-to (w.r.t) the nuclear gene RNase P. Shown are ΔΔct fold change relative to the DMSO control after 3 hrs of treatment with the indicated CCCP doses for both h9-ESCs (**A**) and H9-RGCs (**D**). (**B, E**) Cell viability measurements of H9-ESCs (**B**) and H9-RGCs (**E**) after 24 hrs of treatment with the indicated doses of CCCP. Cell viability was measured using the fluorescence based ApoTox-Glo triplex assay kit and normalized w.r.t DMSO control. (**C, F**) Brightfield images shown are H9-ESCs after 24 hrs of treatment with CCCP (**C**), fluorescence images shown are in the red channel for tdTomato expressing H9-RGCs after 24 hrs of CCCP treatments (**F**). (**G-I**) Mitochondrial level analyzed by the flow cytometry using the mitochondria specific dye MTDR followed by CCCP treatments for 3 hrs. Chart shows the experimental design (**G**), graphs show loss of mitochondria labelled MTDR intensity normalized w.r.t DMSO control at different CCCP doses for H9-ESCs (**H**) and EP1-iPSCs (**I**) compared to the corresponding RGCs. (**J, K**) Cell viability (J) and apoptosis by luminescence-based caspase-3/7 activity (**K**) were measured for EP1-iPSCs using ApoTox-Glo triplex assay kit and normalized w.r.t DMSO control after 24 hrs of treatment with the indicated drugs. Scale bars, 1000 μm (**C**) and 200 μm (**F**). Error bars are SEM. **, p-value < 0.005

### Cell Viability and Apoptosis Measurements

Cell viability and apoptosis were measured using ApoTox-Glo Triplex assay kit (Promega) following manufacturer’s guideline with the CLARIOstar microplate reader (BMG LABTECH). Cell viability was measured by the ratio of fluorescence intensity between 400nm (viability) and 482nm (cytotoxicity) channels, and apoptosis was measured by luminescence-based caspase-3/7 activity. hPSCs (10,000/well) and RGCs (15,000/well) were plated on each well of Matrigel-coated 96well dish in mTeSR containing blebbistatin and N2B27 media respectively. After one day (hPSCs) and three days (RGCs) of recovery, cells were treated with the indicated drugs for 24 hrs and analyzed for cell viability and apoptosis.

### Image Acquisition to Show Cell Viability

Images were acquired after indicated treatments using EVOS FL Imaging System (ThermoFisher Scientific).

### Lysosome/acidic Vesicles Inhibition Assay

20,000 H9-RGCs were plated and grown on Matrigel-coated glass-bottom dish (MatTek) in N2B27 media for three days in the 37°C normoxia incubator (5% CO_2_). After 24 hrs of treatment with the endo-lysosomal inhibitors Baf or HCQ, media was replaced with 100μl of media containing pH sensitive pHrodo-green conjugated dextran solution (ThermoFisher Scientific) (100 μg/ml) for 20min followed by a fresh media exchange after a brief wash. Confocal (Zeiss LSM 710) live images were acquired using the live cell set up with plan-apochromat 40x/oil objective with 1.4 numerical aperture. pHrodo-Green and tdTomato-expressing RGCs were detected with the 488nm and 560nm laser lines respectively.

### Human Cortical Neuron Differentiation

Cortical neurons were differentiated from H1-ESCs as described in Xu et al [37]. Experiments were performed on 100-120 days of post-differentiated cortical neurons.

### Immunofluorescence and Imaging

For measuring ubiquitination level, 20,000 purified RGCs were plated on matrigel-coated glass-bottom dishes (MatTek) for three days followed by 24 hrs of treatment with the indicated drugs. Cells were fixed with 4% paraformaldehyde in PBS for 15min at 37°C followed by 1 hr of blocking at room temperature with blocking solution (PBS with 5% donkey serum and 0.2%Triton X-100). Samples were incubated with primary antibody against ubiquitin (Rabbit-anti-ubiquitin, Cell Signaling Technology, 1:200 dilution) overnight at 4°C. Samples were washed for three times for 5min each with washing solution (PBS with 1% donkey serum and 0.05%Triton X-100) and incubated with the secondary antibody (anti-rabbit-Cy5, 1:500 dilution) in blocking solution for 1 hr at room temperature. Following secondary antibody washed three times, with DAPI added in the second wash.

Cultured human cortical neurons of 100-120 days post-differentiation were immunostained as above with primary antibodies against MAP2 (Mouse-anti-MAP2, Sigma, 1:200 dilution), VGLUT1 (Mouse-anti-VGLUT, SYSY, 1:2500 dilution) and VGAT (Rabbit-anti-VGAT, SYSY, 1:500 dilution). Confocal images were acquired using LSM 710 (Zeiss) as done for pHrodo-Green, but without the live set-up.

### Statistical Analysis

Statistical comparisons between two data sets were done with the Student’s *t*-test. One-way ANOVA tests (Table1) were performed for analysis containing three or more independent groups.

**Table 1.**
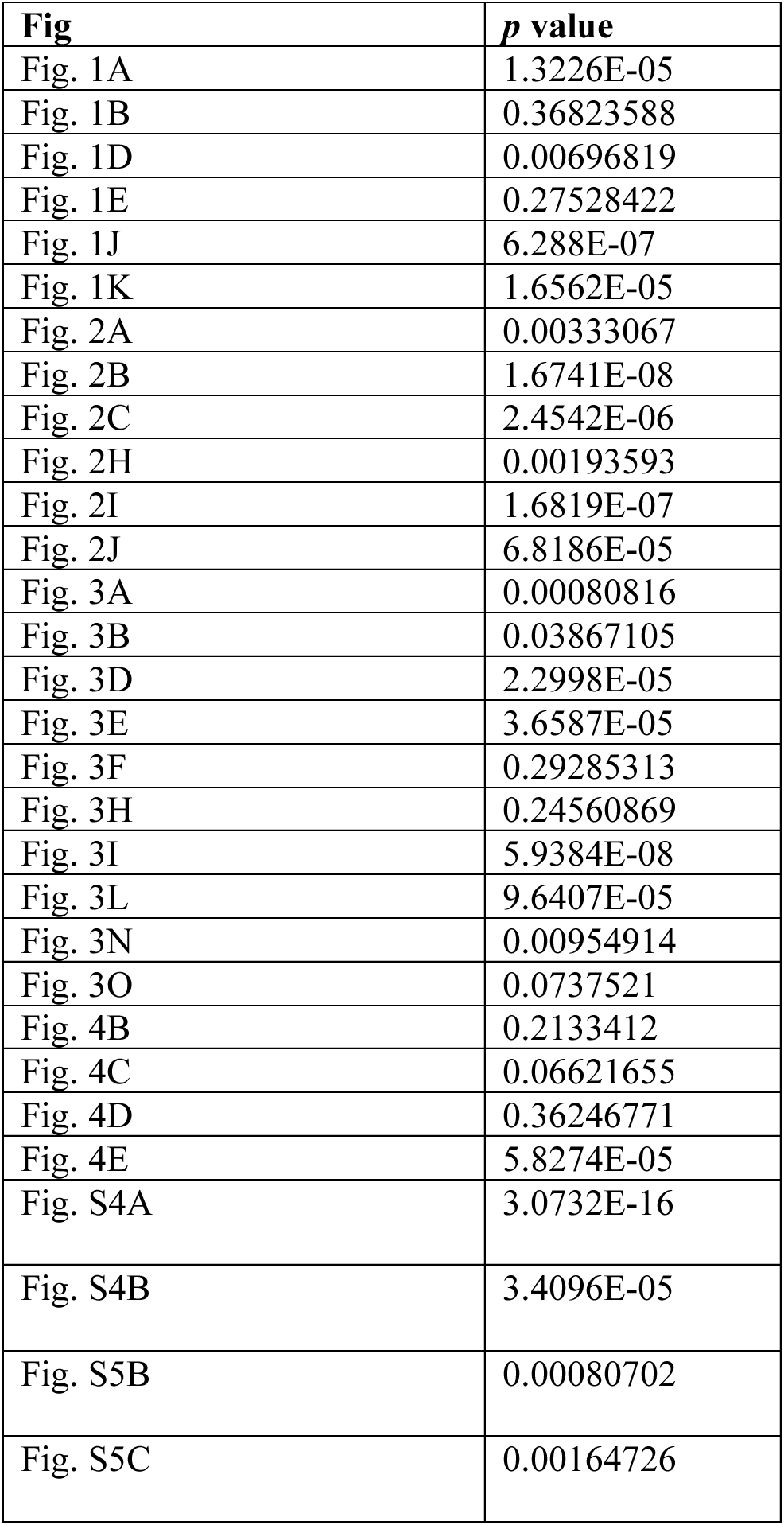
One-way ANOVA test for group of data containing three or more data sets

| **Fig** | ***p* value** |
| --- | --- |
| Fig. 1A | 1.3226E-05 |
| Fig. 1B | 0.36823588 |
| Fig. 1D | 0.00696819 |
| Fig. 1E | 0.27528422 |
| Fig. 1J | 6.288E-07 |
| Fig. 1K | 1.6562E-05 |
| Fig. 2A | 0.00333067 |
| Fig. 2B | 1.6741E-08 |
| Fig. 2C | 2.4542E-06 |
| Fig. 2H | 0.00193593 |
| Fig. 2I | 1.6819E-07 |
| Fig. 2J | 6.8186E-05 |
| Fig. 3A | 0.00080816 |
| Fig. 3B | 0.03867105 |
| Fig. 3D | 2.2998E-05 |
| Fig. 3E | 3.6587E-05 |
| Fig. 3F | 0.29285313 |
| Fig. 3H | 0.24560869 |
| Fig. 3I | 5.9384E-08 |
| Fig. 3L | 9.6407E-05 |
| Fig. 3N | 0.00954914 |
| Fig. 3O | 0.0737521 |
| Fig. 4B | 0.2133412 |
| Fig. 4C | 0.06621655 |
| Fig. 4D | 0.36246771 |
| Fig. 4E | 5.8274E-05 |
| Fig. S4A | 3.0732E-16 |
| Fig. S4B | 3.4096E-05 |
| Fig. S5B | 0.00080702 |
| Fig. S5C | 0.00164726 |

## RESULTS

### RGCs are More Efficient in Degrading Damaged Mitochondria than Their Precursor Stem Cells

To investigate mitochondrial degradation in both human RGCs and in their stem cell origin, we used a CRISPR/Cas9 mediated genetically engineered human embryonic stem cell (hESC-H9) reporter line with a P2A-tdTomato-P2A-Thy1.2 construct introduced into the endogenous RGC-specific POU4F2 (BRN3B) locus [33]. Small molecule-based differentiation followed by immunopurification of Thy1.2-expressing cells yields highly enriched RGCs (Supporting Information Fig. S1) that have been well-characterized transcriptomically and electrophysiologically [33,38]. The reporter line will be referred as “H9-ESCs” and the corresponding RGCs as “H9-RGCs.” To study the effect of mitochondrial damage on RGCs and stem cells, we have used the mitochondrial uncoupler CCCP. Upon mitochondrial damage with CCCP for 3 hrs, H9-ESCs showed no reduction in their mitochondrial level (Fig. 1A), as measured by a qPCR assay that compares the level of mitochondrial gene ND1 DNA to that of nuclear gene RNase P DNA. This result was surprising because CCCP has been reported to induce mitophagy within 1 hr of exposure [39], and hence we expected to see a decrease in mitochondrial content. We hypothesized that due to this apparent lack of appropriate mitochondrial clearance in the H9-ESC cells, which would presumably lead to a buildup of CCCP-induced damaged mitochondria, hence there would be an increase of cell death. However, even with 24 hrs of treatment with CCCP there is no detectable cell death in H9-ESCs (Fig. 1B, C). Contrary to the situation with H9-ESCs, CCCP treatment for 3 hrs reduced the mitochondrial content of H9-RGCs (Fig. 1D). As with the H9-ESCs, CCCP treatment did not result in RGC cell death at 24 hrs (Fig. 1E, F; there was a small decrease in cell viability, but it was not statistically significant). These results suggest that RGCs are may be more efficient in degrading damaged mitochondria than their precursor stem cells.

With the unexpected result of no change in mitochondrial content with CCCP and yet no cell death for H9-ESCs, we next asked whether ESCs might be clearing up damaged mitochondria while simultaneously synthesizing more mitochondria to keep up with their metabolic needs for rapid cell division, and this simultaneous new synthesis might be masking possible mitochondrial degradation. To test this possibility, we tracked mitochondrial levels upon CCCP treatment using the mitochondria-specific dye mitotracker deep red (MTDR) followed by flow cytometry [40] (Supporting Information Fig. S2). MTDR covalently binds to the reduced thiols within the mitochondria matrix proteins and once bound, MTDR remains in the mitochondria independent of mitochondrial membrane potential [41,42]. As CCCP lowers mitochondrial membrane potential and could affect initial MTDR binding, to avoid this potential artifact, mitochondria were labelled with the MTDR dye prior to CCCP treatment, which after appropriate incubation was followed with flow cytometry-based analysis of mitochondrial content (Fig. 1G). With this experimental paradigm, mitochondria synthesized after CCCP treatment will not be labelled, and thus will not be detected, and hence will not mask possible degradation of pre-existing damaged mitochondria. Interestingly, we observed reduced mitochondria levels with increasing doses of CCCP for H9-ESCs (Fig. 1H) and saw similar but more dramatic reduction in mitochondria in H9-RGCs at 5 and 10 μM CCCP (Fig. 1H). To make sure this was not a cell line-specific effect, we performed a parallel experiment with an iPSC-derived POU4F2 reporter line (EP1-iPSCs), again examining both undifferentiated stem cells and differentiated RGCs (EP1-RGCs). In agreement with the H9 results, the iPSCs showed mitochondrial degradation with increasing doses of CCCP and the corresponding RGCs degraded mitochondria more efficiently compared to the undifferentiated iPSCs (Fig. 1I). Of note, EP1-iPSCs showed relatively reduced mitochondrial clearance compared to the H9-ESCs (Fig. 1H, I) and upon CCCP damage, correspondingly showed reduced cell viability and increased apoptosis (Fig. 1J, K), further supporting the hypothesis that inefficient degradation of damaged mitochondria may lead to apoptotic cell death. These results suggest that human RGCs more efficiently remove damaged mitochondria than their precursor stem cells, which may play a key role in the long-term survival of RGCS in humans. Investigating the pathways involved in degrading damaged mitochondria in hRGCs could be therapeutically important as modulation of the pathways involved could potentially be used to enhance hRGC survival.

### The Endo-lysosomal Pathway is Required for hRGCs but not for hESCs to Degrade Damaged Mitochondria

To better define possible cell type-specific mechanisms of mitochondrial quality control in hRGCs, we first tested the role of the endo-lysosomal pathway in degrading damaged mitochondria in hPSCs and hRGCs. Mitochondrial levels were measured after CCCP treatment both in the presence and absence of the endo-lysosomal inhibitors hydroxychloroquine (HCQ) [43] and Bafilomycin A1 (Baf) [44]. qPCR-based analysis showed that individual treatment with CCCP, HCQ, or Baf, as well as CCCP with Baf, did not affect mitochondrial level in H9-ESCs (Fig. 2A). Tracking MTDR labelled mitochondrial content upon CCCP treatment showed mitochondrial degradation in hPSCs (Fig. 1H, I). Our inability to detect an increase in mitochondria levels upon endo-lysosomal inhibition may suggest the existence of alternative pathway in H9-ESCs for degrading damaged mitochondria. We observed significant cell death and apoptosis when cells were treated with Baf but did not observe similar cell death with HCQ treatment (Fig. 2B, C). This could be due to the requirement of endo-lysosomal activity and autophagy pathway for other cellular functions, such as non-mitochondrial protein and organelle homeostasis [45,46]. The differential effects of Baf and HCQ on H9-ESC survival could be due to the distinct modes of action of the two inhibitors [47]. We next tested if inhibition of the mitochondrial electron transport chain (mETC) with oligomycin-antimycin (OA) would cause mitochondrial degradation in ESCs. Interestingly, we did not observe reduced mitochondrial levels with OA treatment; on the contrary, we observed increased mitochondrial levels when cells were treated alone or in combination with Baf (Fig. 2A). This finding is consistent with a prior report that inhibition of the mETC is associated with the inhibition of autophagy [48], which could account for the observed increase in mitochondrial content, and the increased cell death and apoptosis with OA treatment (Fig. 2B, C). Since oligomycin has been reported to increase inner mitochondria membrane potential (ΔΨ_m_) [49,50], we further asked if increasing ΔΨ_m_ by inhibition of the mitochondrial permeability transition pore (mPTP) with cyclosporin A (CsA) [50] could also block mitochondrial degradation. In agreement with the OA result, we observed increased mitochondrial levels in H9-ESCs treated with CsA (Fig. 2D, E). Presumably because inhibition of damaged mitochondrial degradation can be toxic, CsA treatment also caused increased cell death and activation of apoptosis (Fig. 2F, G).

**Figure 2.**
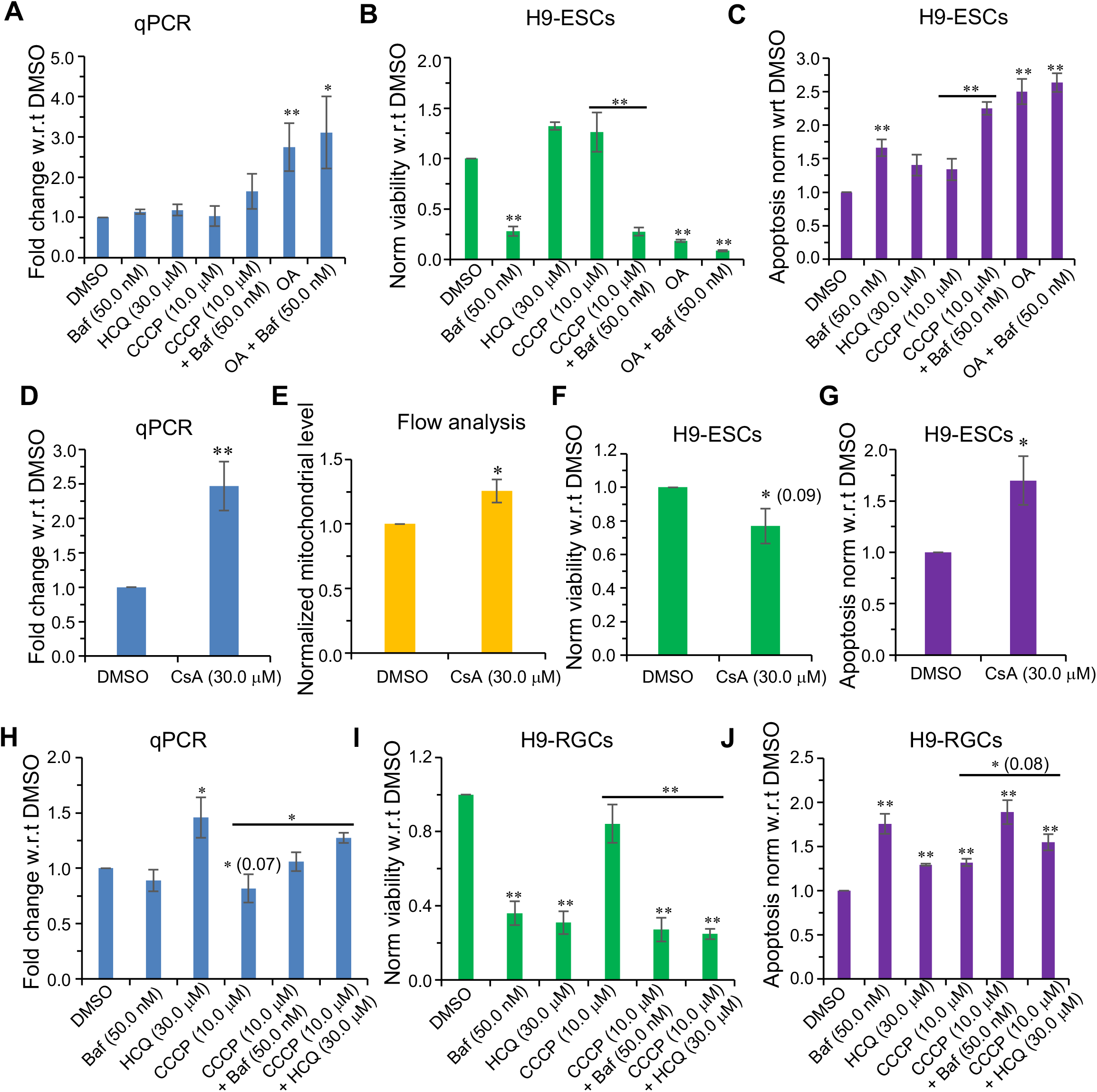
hRGCs but not hESCs predominantly use endo-lysosomal pathway for degrading mitochondria. (**A, D, H**) qPCR-based analysis of the mitochondrial content for H9-ESCs (**A, D**) and H9-RGCs (**H**) after 24 hrs of treatment with the indicated drugs, quantification and the analysis are done as in (Figure. 1). (**B, F, I**) Shown are cell viability measurements after 24 hrs of treatment with the indicated drugs for H9-ESCs (**B, F**) and H9-RGCs (**I**) are done as in (Figure. 1). (**C, G, J**) Quantifications represent cellular apoptosis, measured by luminescence-based caspase-3/7 activity for H9-ESCs (**C, G**) and H9-RGCs (**J**). (**E**) Flow cytometry-based analysis of the MTDR labelled mitochondria for H9-ESCs after 24 hrs of treatment with the indicated drug, quantification shows normalized average intensity in the MTDR channel w.r.t DMSO. Error bars are SEM. **, p-value < 0.01; *, p-value < 0.05.

To test whether H9-RGCs degrade damaged mitochondria via endo-lysosomes, we blocked endo-lysosomal activity with Baf and HCQ. HCQ (with and without CCCP), but not Baf, increased mitochondrial content (Fig. 2H), indicating that HCQ is a more potent inhibitor of mitophagy in H9-RGCs than in hESCs. As expected, presumably due to its inhibition of mitochondrial clearance, HCQ caused hRGC death and apoptosis (Fig. 2I, J). Although we did not observe increased levels of mitochondria with Baf treatment (Fig. 2H), we did observe increased H9-RGC apoptosis and cell death (Fig. 2I, J). As an indication that they were having their expected pharmacological activities, both Baf and HCQ increased the pH of the acidic endo-lysosomal vesicles as shown by a decrease in fluorescence dots of the pH-sensitive dye pHrodo-green dextran (Supporting Information Fig. S3). The explanation of the differences between the effects of Baf and HCQ is unclear, but may reflect that the two drugs differentially affect the endo-lysosomal compartments which have been reported [47].

The above data suggest differentiated RGCs are different from their origin stem cells in terms of using endo-lysosomes for degrading damaged mitochondria. For stem cells, inhibition of endo-lysosomes was toxic but did not increase mitochondrial content. While in H9-RGCs inhibition of endo-lysosomal pathway both inhibited mitochondrial degradation as well as reduced RGC survival, suggesting RGCs predominantly use endo-lysosomal pathway for mitophagy and cellular homeostasis. Choice for using endo-lysosomal pathway versus UPS to maintain healthy cellular homeostasis is critical for cell survival. With the apparent difference between hESCs and hRGCs in choosing endo-lysosomal pathway and the potential involvement of UPS in neurodegenerative diseases [51], led us to ask the role of UPS for mitochondrial clearance and neuro-protection in hRGCs and for the origin stem cells.

### The UPS is Required for Mitochondrial Degradation and Cell Survival for hPSCs but not for hRGCs

As an alternative to the endo-lysosomal pathway, the proteasomal (UPS) pathway is the other major cellular quality control pathway for the protein and organelle homeostasis [52]. We next investigated the role of UPS in mitochondrial clearance by using the drug bortezomib to inhibit the proteasome’s 20S core particle [53]. Unexpectedly, we found that inhibiting proteasome in H9-ESCs increased mitochondrial levels in a dose dependent manner (Fig. 3A, B). As accumulation of damaged mitochondria could lead to cellular toxicity, we further observed cell death and activation of the apoptotic pathway in H9-ESCs with bortezomib treatment (Fig. 3C-E). To test if the observed effect was specific to the ESC line, we also inhibited proteasome function in EP1-iPS cells and also observed dose dependent cell death with concomitant activation of apoptosis (Supporting Information Fig. S4). These data suggest that proteasomal activity is critical for basal level mitochondrial clearance and survival of hPSCs. We next asked if proteasomal activity is required for the clearance of acutely damaged mitochondria in H9-ESCs. To test this, we induced mitochondrial damage by CCCP both in presence and the absence of bortezomib. In agreement with our hypothesis, we observed an increase level of mitochondria when proteasomal clearance was blocked by bortezomib compared to CCCP alone (Fig. 3F), suggesting that hPSCs use predominantly the proteasomal pathway to degrade damaged mitochondria.

**Figure. 3.**
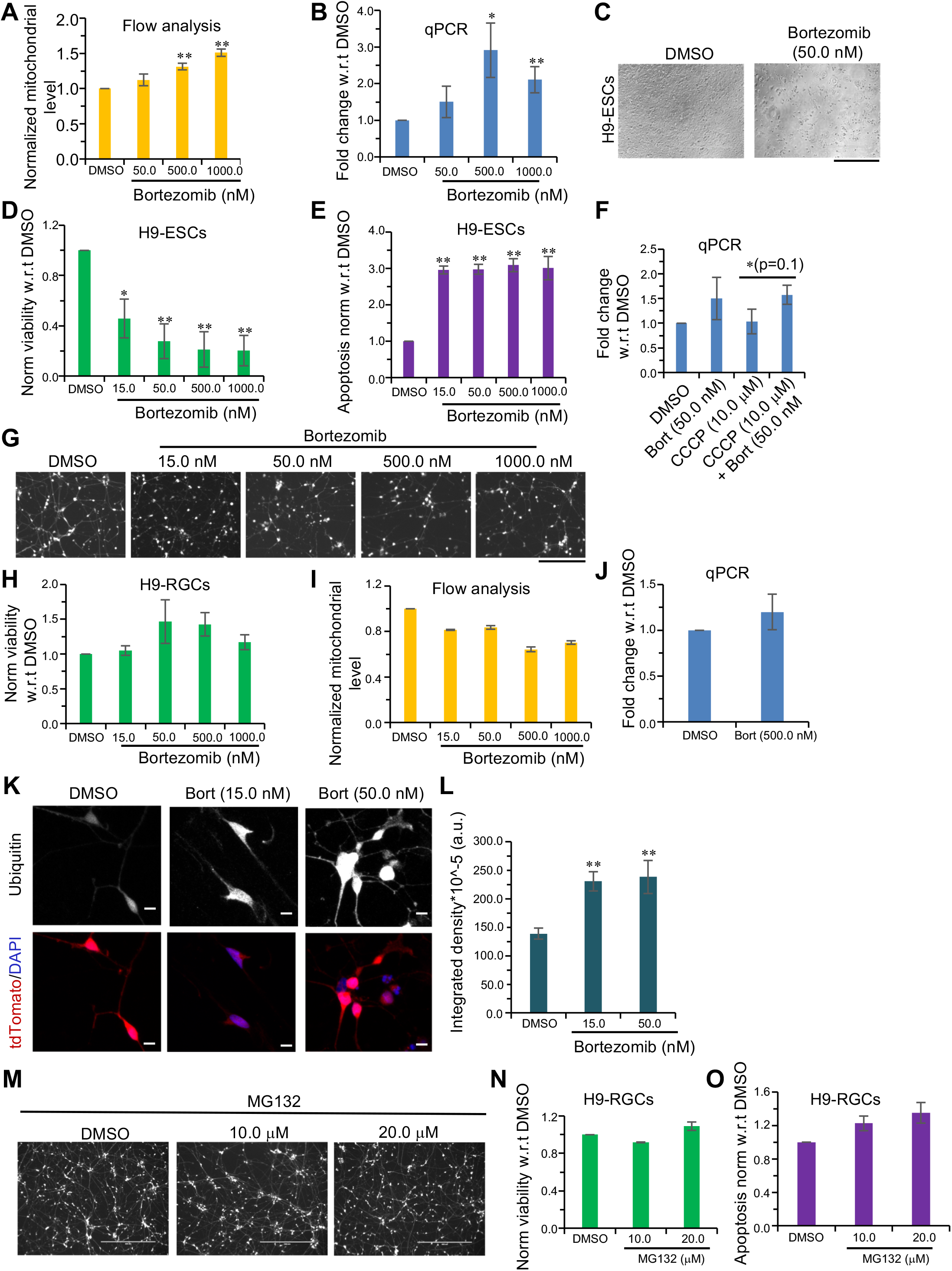
hESCs but not hRGCs predominantly use proteasomal pathway for degrading mitochondria. (**A, I**) Flow cytometry-based analysis of the MTDR labelled mitochondria in H9-ESCs (**A**) and H9-RGCs (**I**) after 24 hrs of treatment with the indicated doses of bortezomib. (**B, F, J**) qPCR analysis of the mitochondrial content in H9-ESCs (**B, F**) and H9-RGCs (**J**) after 24 hrs of treatment with the indicated drugs, quantification and the analysis are done as in (Figure. 1). (**C**) Brightfield images showing cell death in the H9-ESCs after 24 hrs of treatment with the indicated bortezomib (Bort) dose. (**D, H, N**) Shown are cell viability measurements after 24 hrs of treatment with the indicate drugs for H9-ESCs (**D**) and H9-RGCs (**H, N**) as done in (Figure 1). (**E, O**) Quantifications represent cellular apoptosis, measured by luminescence-based caspase-3/7 activity for H9-ESCs (**E**) and H9-RGCs (**O**). (**G, M**) Fluorescence images shown are in the red channel for the tdTomato expressing H9-RGCs after 24 hrs of bortezomib (Bort) (**G**) and MG132 (**M**) treatments with the indicated doses. (**K, L**) Images shown are the sum projections of the confocal z-stacks on immunofluorescence against ubiquitin in H9-RGCs after 24 hrs of treatment with the indicated bortezomib doses (**K**), and quantification shows the integrated fluorescence intensity from the sum-projections of individual cell (**L**). Scale bars, 400 μm (**C, M**), 200 μm (**G**) and 10 μm (**K**). Error bars are SEM. **, p-value < 0.01; *, p-value < 0.05.

To investigate if proteasomal activity is required for mitochondrial degradation and hRGC survival, we treated H9-RGCs with different doses of bortezomib for 24 hrs followed by viability and mitochondrial level measurements. Interestingly, unlike hPSCs, inhibiting proteasomal activity did not affect the survival of H9-RGCs (Fig. 3G, H). Next, we tested whether inhibiting the UPS with bortezomib affects mitochondrial homeostasis in H9-RGCs. Contrary to the hPSCs, proteasomal inhibition did not increase the mitochondrial level in H9-RGCs (Fig. 3I, J). To test the generality of these observations, proteasomal inhibition was further tested on iPSC derived RGCs (EP1-RGCs). EP1-RGCs showed mild cell death effect with a moderate increase in apoptotic activity with bortezomib treatment (Supporting Information Fig. S5A-C), but to a considerably lesser extent than observed with EP1-iPSCs, especially with respect to cell death (Supporting Information Fig. S4). With the observed differences in proteasomal regulation between hPSCs and hRGCs, we next asked if the UPS is still active in the hRGCs. Ubiquitinated proteins and organelles are degraded through the proteasome [54,55], hence inhibiting UPS activity should increase the ubiquitinated protein level. In support of our hypothesis, we found a bortezomib dose-dependent increase in the ubiquitinated protein level in H9-RGCs (Fig. 3K, L). We further tested the bortezomib results using another very potent proteasome inhibitor, MG132 [53,56]. In agreement with the bortezomib data, MG132 also did not induce cell death or apoptosis in H9-RGCs (Fig. 3M-O).

These results suggest a switch in the mitochondrial degradation pathways from the proteasome to the endo-lysosomal pathway during human RGC differentiation, making the lysosomal-autophagy pathway a potential therapeutic target for improving mitochondrial health and therefore hRGC survival in glaucoma and in other forms of optic neuropathy patients. While these findings could be important for improving hRGC health, we were additionally interested to see if these phenomena are specific for hRGCs or also true for other types of human neurons. To address this question, we have additionally differentiated human cortical neurons and tested the effect of endo-lysosomal and proteasomal inhibition on them.

### Human Cortical Neurons are Susceptible to the Endo-lysosomal Inhibition but not to the Proteasome Inhibition

To study the role of proteasomal and endo-lysosomal pathways for cortical neuron survival, we differentiated cortical neurons from human stem cells (H1-ESCs) following published methods [57]. Cultured human cortical neurons were tested and shown to be positive for expression of the mature neuronal marker microtubule-associated protein 2 (MAP2), inhibitory marker vesicular GABA transporter (VGAT), and excitatory marker vesicular glutamate transporter (VGLUT) (Fig. 4A). To test the effect of endo-lysosomal inhibition, cells were treated with HCQ. Similar to hRGCs, we observed significant cell death and corresponding activation of apoptosis (Fig. 4B, C), suggesting endo-lysosomal pathway is important for cellular homeostasis in cortical neurons.

**Figure. 4.**
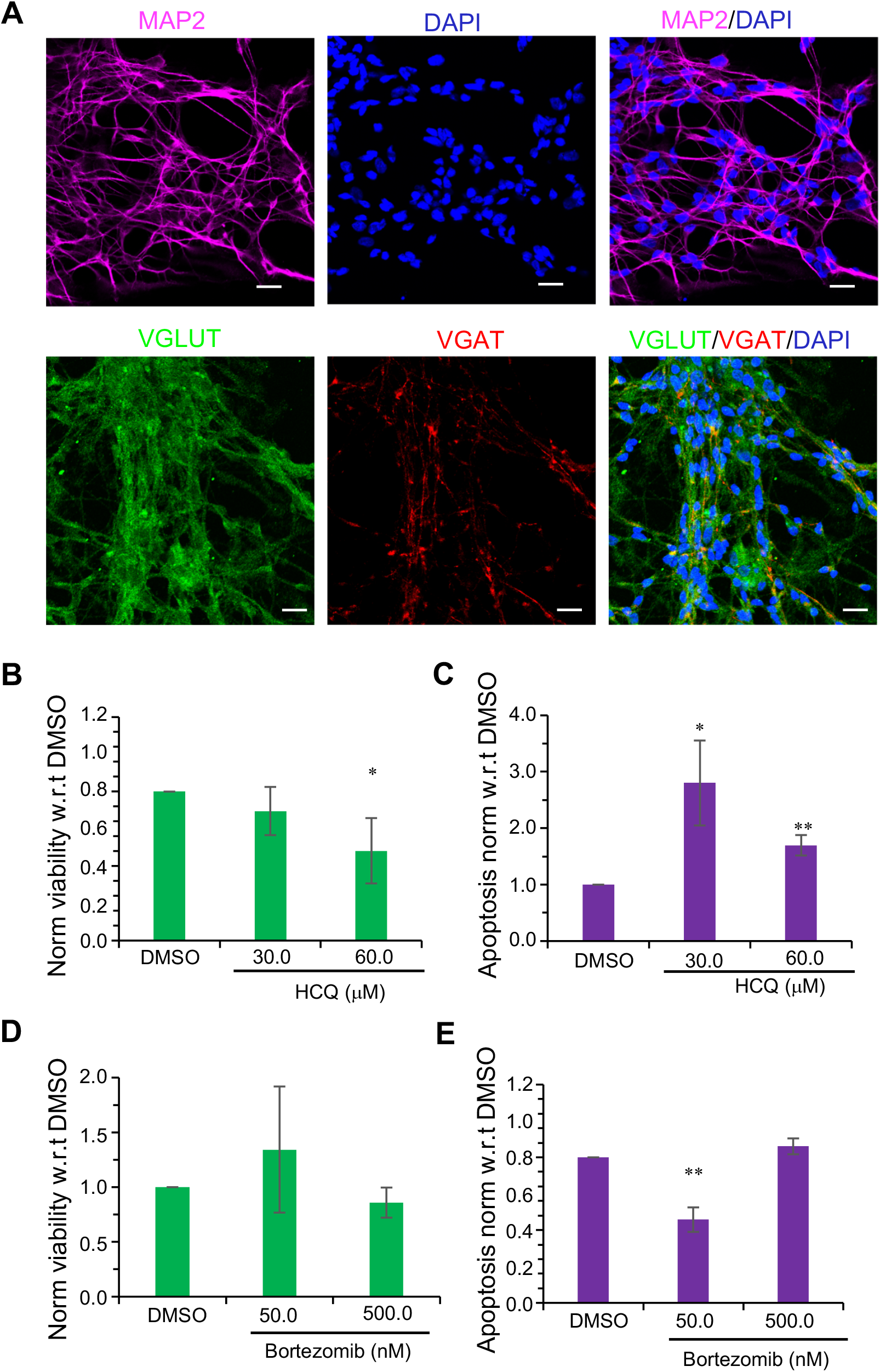
Effect of UPS and endo-lysosomal pathway inhibition on human cortical neuron survival. (**A**) Shown are the confocal images of immunofluorescence against neuronal marker MAP2, excitatory marker VGLUT and the inhibitory marker VGAT. (**B-E**) After 24 hrs of treatment with the indicated drugs, cell viability (**B, D**) and apoptosis (**C, E**) were measured using ApoTox-Glo triplex assay kit. Scale bars are 20 μm. Error bars are SEM. **, p-value < 0.01; *, p-value < 0.05.

Next, we tested the effect of proteasomal inhibition by treating cortical neurons with the proteasome inhibitor bortezomib. Interestingly, like hRGCs, proteasome inhibition did not cause cell death for human cortical neurons (Fig. 4D, E).

Taking together, our data suggest endo-lysosomal pathway may be the predominant pathway for degrading damaged mitochondria and maintaining cellular homeostasis for not only human RGCs but also for cortical neurons. It will be interesting to test the cellular preference in choosing among proteasomal and the endo-lysosomal pathways in other differentiated human cell types as well.

## DISCUSSION

Our study identifies pathways important for maintaining healthy mitochondrial homeostasis in human RGCs, which is a step forward for developing strategies to enhance RGC viability under disease conditions. The results shown here suggest three key points: first, human RGCs are more efficient in clearing up damaged mitochondria than their precursor stem cells; second, the proteasomal pathway is essential for stem cell survival but not for RGCs; and third, during RGC maturation from stem cells, the pathway for mitochondrial clearance shifts from the proteasomal to the endo-lysosomal pathway.

While we observed mitochondrial degradation in stem cells is dependent on UPS, a question still remains on how proteasomes degrade mitochondria? An elegant study by Chan et al (2011) suggests that this could happen by Parkin mediated activation of the UPS. Upon its translocation to mitochondria, Parkin activates the 26S proteasome, leading to the degradation of the mitochondrial outer membrane proteins. For hPSCs, a similar mechanism may lead to UPS mediated mitochondrial degradation as well. However, our data suggest UPS dependent mitochondrial degradation could depend on the mitochondrial inner membrane potential (ΔΨ_m_). We have seen that when ESCs were treated with OA or CsA, which are known to increase ΔΨ_m_ [49,50], mitochondrial degradation was inhibited (Fig. 2A, D, E). However, when treated with the uncoupler CCCP, which abolishes ΔΨ_m_ [49], mitochondrial degradation was induced (Fig. 1H, I). This suggests ΔΨ_m_ could negatively regulate UPS mediated mitochondrial degradation. Further efforts will require to understand the mechanism of how ΔΨ_m_ can regulate UPS mediated mitochondrial degradation. Even though we observed that inhibition of endo-lysosomes did not block mitophagy in ESCs, endo-lysosome involvement could not be ruled out as a previous report had shown inhibiting lysosomes increased mitochondrial content in hematopoietic stem cells (HSCs) [59].

Our study indicates that the endo-lysosomal pathway is the primary route for degrading damaged mitochondria in hRGCs. This is significant as it makes the endo-lysosomal pathway a potential therapeutic target to enhance mitochondrial homeostasis to increase hRGC survival. Furthermore, genetic analyses has identified the mutation in the mitophagy adaptor protein Optineurin (Optn) in normal tension glaucoma (NTG) patients [31], which makes our finding even more therapeutically relevant. Mutations in the MQC pathway proteins Mitofusin1/2 (Mfn1/2) [60] and mitochondrial DNA mutations are also associated with other forms of optic neuropathy such as LHON and neuropathy, ataxia and retinitis pigmentosa (NARP) [61]. A cellular-level intervention to maintain healthy mitochondria and mitigate optic nerve disease progression will be aided by increased understanding of the damaged mitochondrial clearance pathways in human RGCs.

## CONCLUSION

A switch from the proteasome to the endo-lysosomal pathway for mitochondrial degradation and cell survival during RGC differentiation is significant, as this could be a general shift for other proteins and organelle homeostasis as well. The impact of such a developmental process could be twofold: first, UPS-mediated protein degradation is an ATP-dependent process and hence requires energy [62], so avoiding the UPS for protein and organelle homeostasis could be a big energy saving strategy for highly energy-dependent RGCs [63]. Second, endo-lysosomal pathway being the primary mitochondria degradation pathway for RGCs makes this pathway a therapeutic target for RGC protection in mitochondria-based optic neuropathies. Additionally, enhancing the endo-lysosome pathway could also be a valid approach for other neurodegenerative diseases since our study indicates that differentiated human cortical neurons also use the endo-lysosomal pathway for their survival.

## ACKNOWLEDGMENTS

We thank Drs. Jin-Chong Xu, Ted M. Dawson and Valina L. Dawson for providing differentiated human cortical neurons. We also thank Drs. Sayantan Datta and James Tahara Handa for helping us to develop qPCR based mitochondrial measurements. This work was supported by grants from the NIH (P30 EY001765, K99 EY028223, and R01 EY026471), Research to Prevent Blindness and generous gifts from the Guerrieri Family Foundation.

## DISCLOSURE OF POTENTIAL CONFLICTS OF INTRESET

The authors declare no potential conflicts of interest.

## DATA AVAILABILITY STATEMENT

The data used in the current study are available from the corresponding authors upon reasonable request.

## Author Contributions

Arupratan Das: Conception and design, financial support, collection and assembly of data, data analysis and interpretation, manuscript writing, final approval of manuscript; Claire M. Bell: collection and assembly of data; Cynthia A. Berlinicke: collection and assembly of data, manuscript writing; Nicholas Marsh-Armstrong: data analysis and interpretation, manuscript writing; Donald J. Zack: conception and design, financial support, administrative support, collection and assembly of data, data analysis and interpretation, manuscript writing, final approval of manuscript.

---

**Supplementary Fig. S1.**
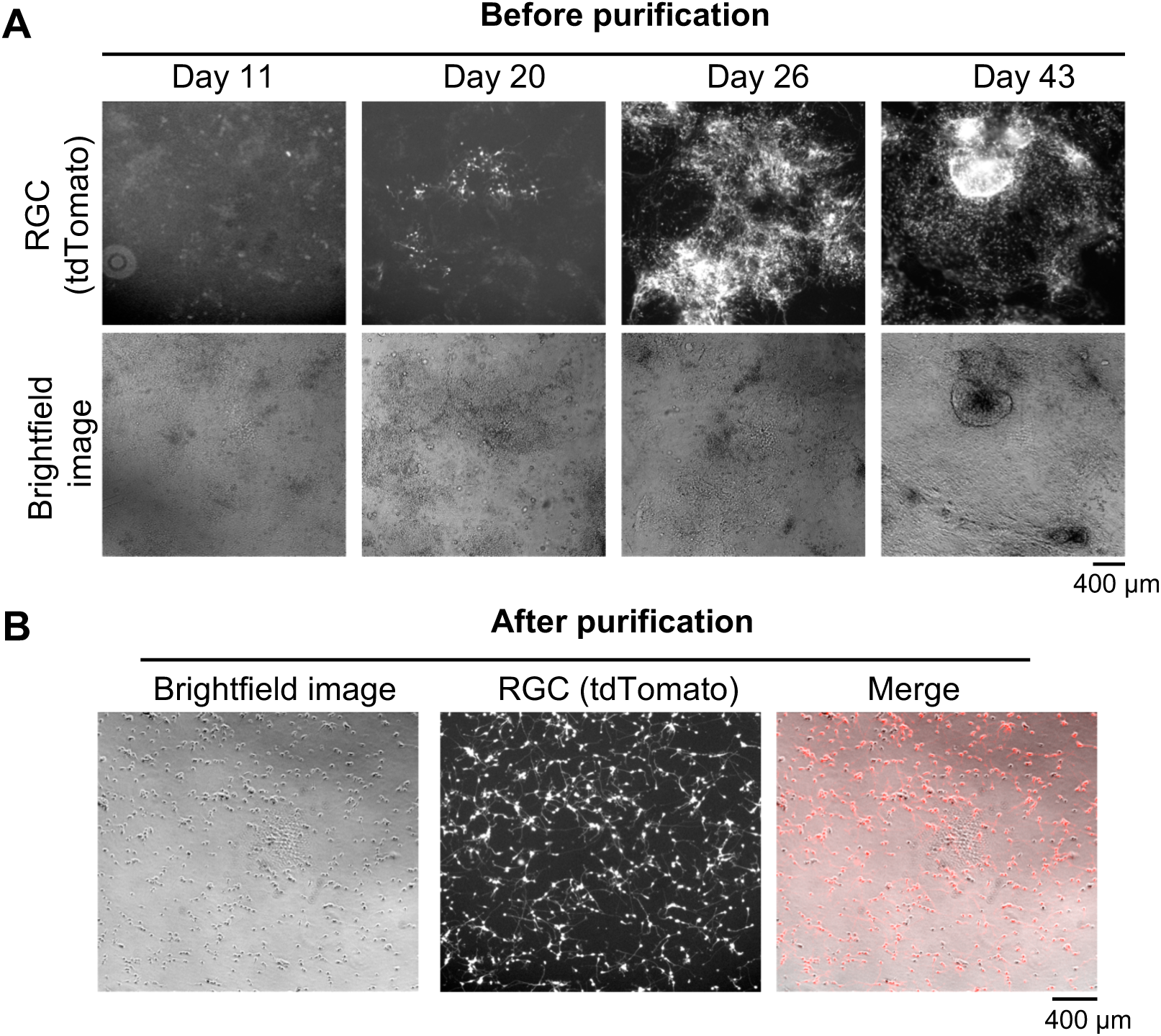
hRGC differentiation from the H9-ESCs. (**A**) Images shown are the representatives of different time points for RGC differentiation, tdTomato positive cells at day 20, 26 and 43 indicates successful RGC differentiation. (**B**) Images shown are the H9-RGCs immunopurified against the surface antigen Thy1.2 and grown on matrigel coated tissue culture dish, high overlap between the brightfield and the tdTomato channel indicates highly pure RGC culture. Scale bars, 400 μm.

**Supplementary Fig. S2.**
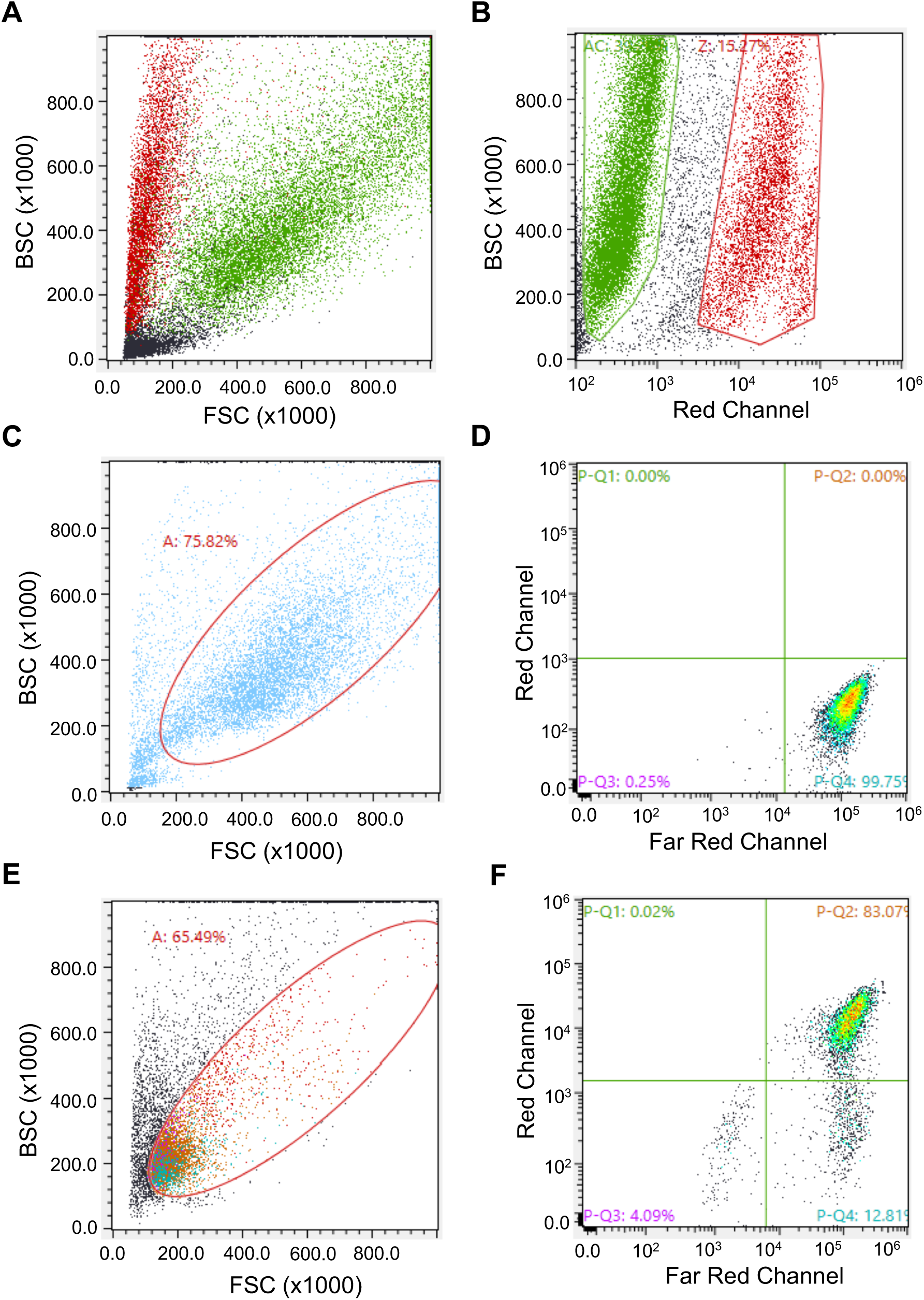
Flow cytometry-based analysis of the mitochondrial content. (**A**) Forward-backward scatter plot of the H9-ESCs labelled with the dead cell dye propidium iodide (PI). (**B**) Diagonally distributed green dots in ‘**A**’ are low in PI intensity representing live cells and red dots along the BSC axis in ‘**A**’ are high in PI intensity representing dead cells. This allowed empirically to select diagonally distributed live cell population for analysis. (**C**) Diagonally distributed live H9-ESCs were gated (red oval) for analysis. (**D**) Live H9-ESCs labelled with mitochondria dye MTDR (far-red) as shown in P-Q4 quadrant were analyzed for average MTDR intensity. (**E**) Diagonally distributed live H9-RGCs were gated (red oval) for analysis. (**F**) Live H9-RGCs positive for both tdTomato (red) and MTDR (far-red) distributed in the P-Q2 quadrant were analyzed for average MTDR intensity.

**Supplementary Fig. S3.**
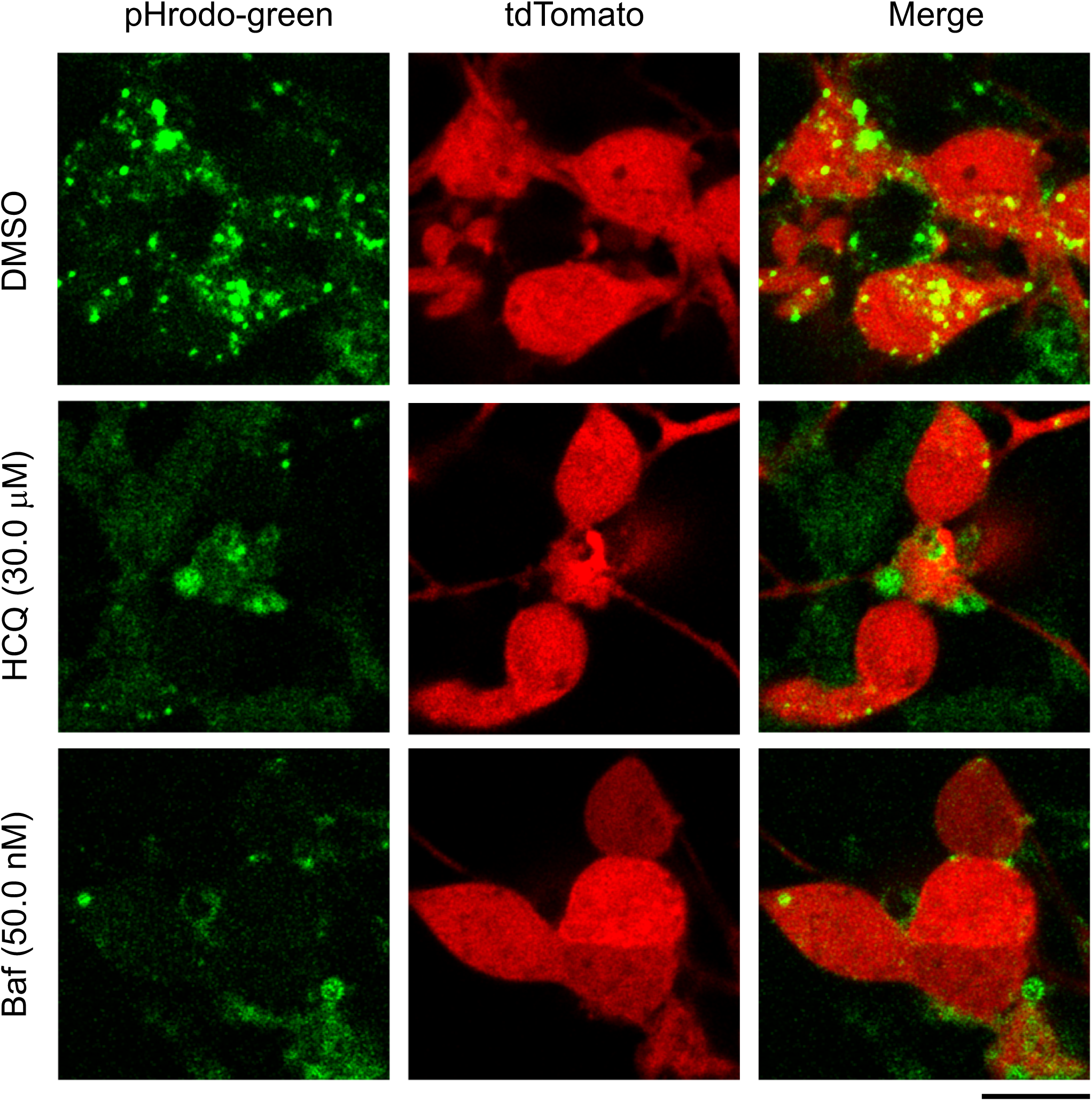
Bafilomycin A1 (Baf) and hydroxychloroquine (HCQ) increased pH in RGCs. Confocal images shown are live H9-RGCs after 24 hrs of treatment with the indicated drugs followed by 20min incubation with the pH sensitive pHrodo-green conjugated dextran. Scale bar, 10 μm.

**Supplementary Fig. S4.**
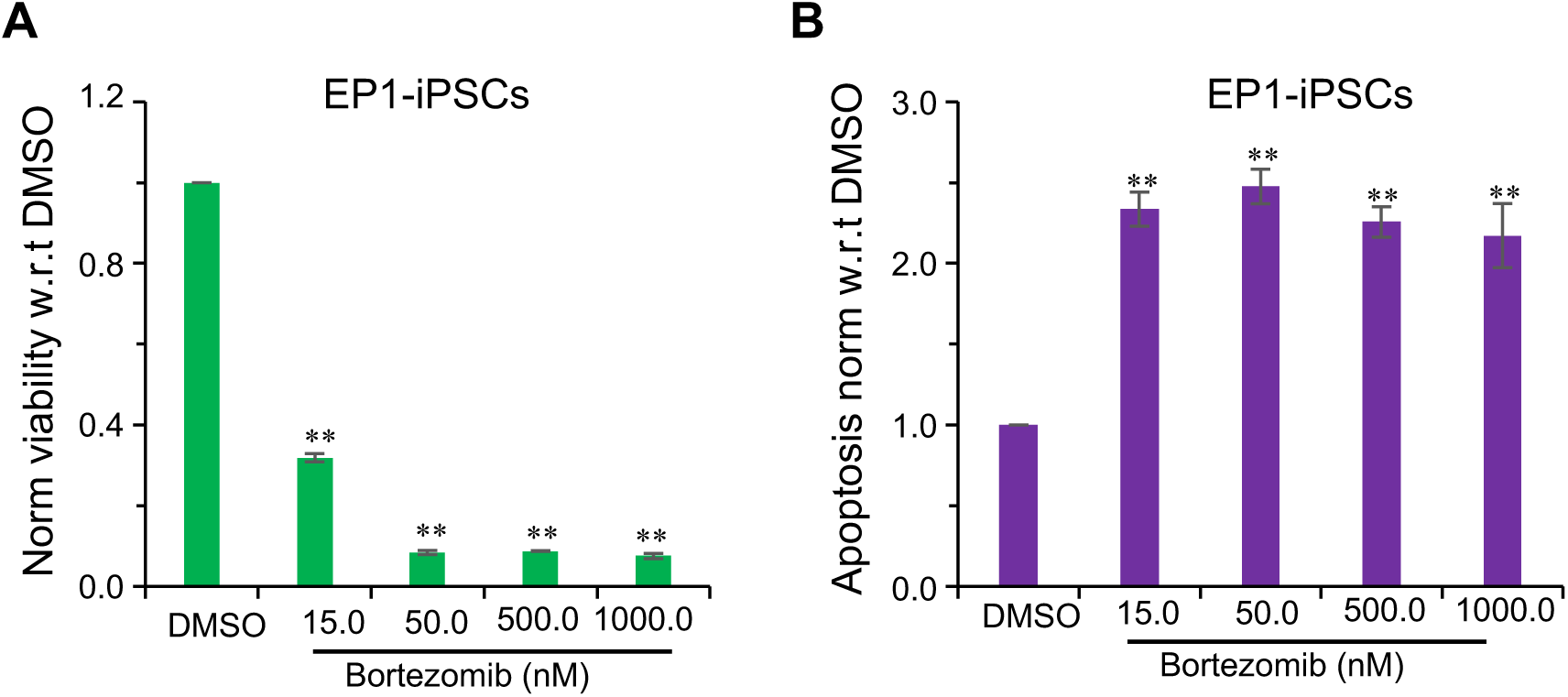
Proteasomal activity is required for the iPSC survival. (**A, B**) Cell viability (**A**) and caspase-3/7 activity for apoptosis (**B**) in EP1-iPSCs were measured using ApoTox-Glo triplex assay after 24 hrs of treatment with the bortezomib at the indicated doses. Error bars are SEM. **, p-value < 0.005.

**Supplementary Fig. S5.**
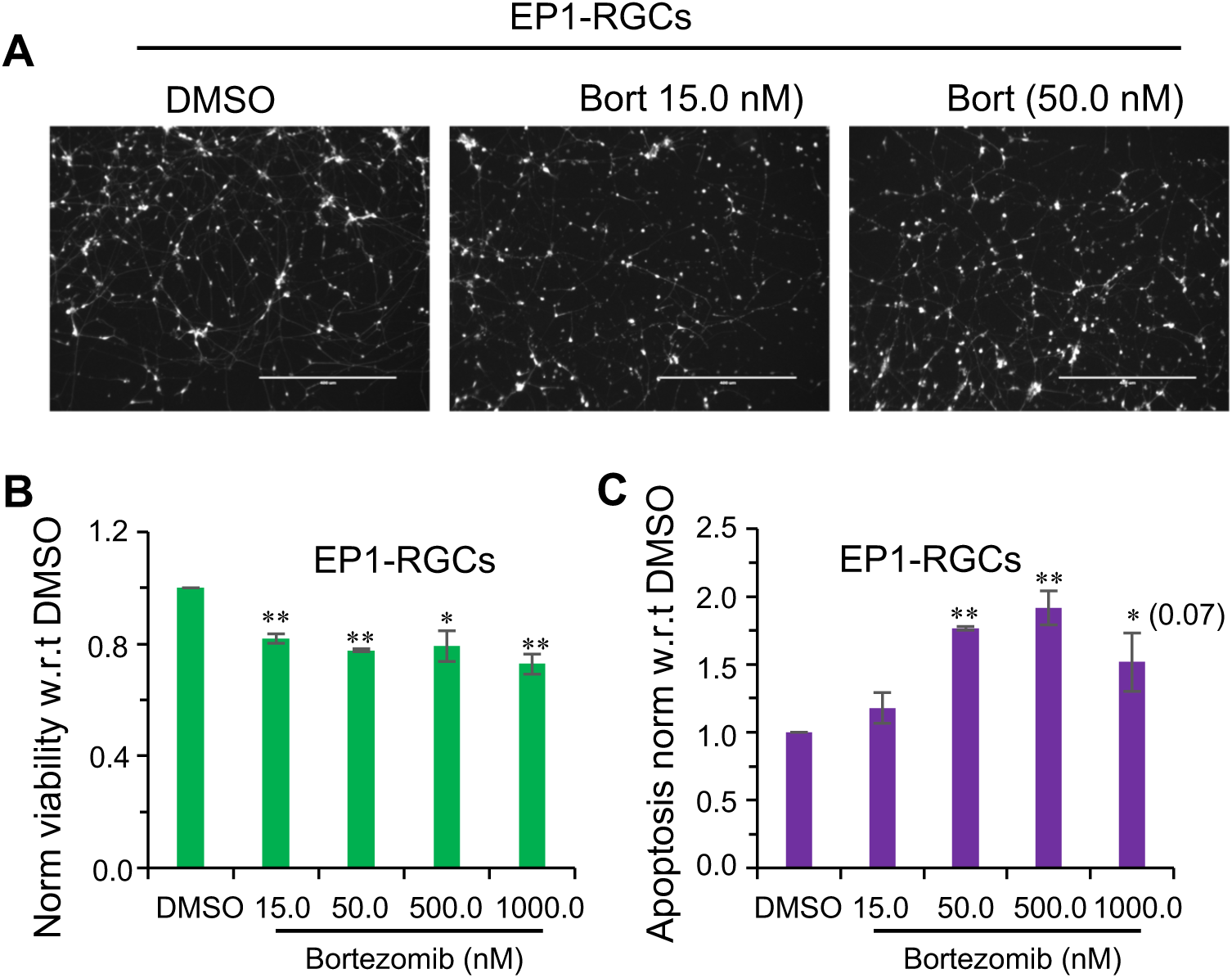
Effect of proteasomal inhibition on iPSC derived RGCs. (**A**) Images shown are the tdTomato expressing EP1-RGCs after 24 hrs of treatment with the bortezomib at the indicated doses. (**B, C**) Cell viability (**B**) and caspase-3/7 activity for apoptosis (**C**) were measured using ApoTox-Glo triplex assay after 24 hrs of treatment with bortezomib at the indicated doses. Scale bars, 400 μm. Error bars are SEM. **, p-value < 0.01; *, p-value < 0.05

